# Comprehensive analysis of clustered mutations in cancer reveals recurrent APOBEC3 mutagenesis of ecDNA

**DOI:** 10.1101/2021.05.27.445689

**Authors:** Erik N. Bergstrom, Jens Luebeck, Mia Petljak, Vineet Bafna, Paul S. Mischel, Reuben S. Harris, Ludmil B. Alexandrov

## Abstract

Clustered somatic mutations are common in cancer genomes with prior analyses revealing several types of clustered single-base substitutions, including doublet- and multi-base substitutions, diffuse hypermutation termed *omikli*, and longer strand-coordinated events termed *kataegis*. Here, we provide a comprehensive characterization of clustered substitutions and clustered small insertions and deletions (indels) across 2,583 whole-genome sequenced cancers from 30 cancer types. While only 3.7% of substitutions and 0.9% of indels were found to be clustered, they contributed 8.4% and 6.9% of substitution and indel drivers, respectively. Multiple distinct mutational processes gave rise to clustered indels including signatures enriched in tobacco smokers and homologous-recombination deficient cancers. Doublet-base substitutions were caused by at least 12 mutational processes, while the majority of multi-base substitutions were generated by either tobacco smoking or exposure to ultraviolet light. *Omikli* events, previously attributed to the activity of APOBEC3 deaminases, accounted for a large proportion of clustered substitutions. However, only 16.2% of *omikli* matched APOBEC3 patterns with experimental validation confirming additional mutational processes giving rise to *omikli. Kataegis* was generated by multiple mutational processes with 76.1% of all *kataegic* events exhibiting AID/APOBEC3-associated mutational patterns. Co-occurrence of APOBEC3 *kataegis* and extrachromosomal-DNA (ecDNA) was observed in 31% of samples with ecDNA. Multiple distinct APOBEC3 *kataegic* events were observed on most mutated ecDNA. ecDNA containing known cancer genes exhibited both positive selection and *kataegic* hypermutation. Our results reveal the diversity of clustered mutational processes in human cancer and the role of APOBEC3 in recurrently mutating and fueling the evolution of ecDNA.

## INTRODUCTION

The genomes of cancer cells harbor somatic mutations imprinted by the activities of different mutational processes^1,2^. Most single-base substitutions and small insertions and deletions (indels) are independently scattered across the genomic landscape; however, a subset of substitutions and indels tend to cluster together^3,4^. This clustering has been attributed to a combination of heterogeneous mutation rates across the genomic landscape, biophysical characteristics of exogenous carcinogens, dysregulation of endogenous processes, and the occurrence of larger events associated with genome instability; amongst others^4-16^. Prior analyses of clustered mutations have focused on single-base substitutions and revealed several classes of clustered events, including doublet- and multi-base substitutions^1,9,16-19^, diffuse hypermutation termed *omikli*^15^, and longer events termed *kataegis*^12,14,16,20^. The majority of *kataegic* events were found to be strand-coordinated which is usually defined as sharing the same strand and reference allele^2,16^. To the best of our knowledge, analysis of cluster indels has never been performed.

Doublet-base substitutions have been extensively examined revealing multiple endogenous and exogenous mutational processes that can cause these events, including, failure of DNA repair pathways and exposure to environmental mutagens^1,2,16,17^. In contrast, multi-base substitutions have not been comprehensively explored presumably due to their small numbers in most cancer genomes. Moreover, only a handful of reported processes have been associated with *omikli* and *kataegic* events with majority of these processes attributed to AID/APOBEC3 family of deaminases^4,5,12,14-16,21-24^. For example, in B-cell lymphomas clustered tracks of C>T and C>G mutations at WR<u>C</u>Y motifs are the result of direct replication over AID lesions^21^. Alternatively, AID-induced lesions can be processed by the mismatch repair pathway that recruits the error-prone DNA polymerase η resulting in non-canonical AID mutations^21^. In addition to AID, the APOBEC3 enzymes, which are typically responsible for anti-viral responses and for limiting the mobility of mobile elements^25-31^, are a substantial contributor of clustered mutational events^2,4,12,14-16,24,32^. Specifically, the APOBEC3 enzymes give rise to *omikli* and *kataegis* by requiring single-stranded DNA as a substrate^14,15,24,32^. *Omikli* were found enriched in early replicating regions and more prevalent in microsatellite stable tumors indicating a role of mismatch repair in exposing short single-stranded DNA regions while processing mismatched bases during replication^15^. Further, the differential activity of mismatch repair towards gene-rich regions results in an increased mutational burden of *omikli* mutations within cancer driver genes^15^. *Kataegis* is less prevalent than *omikli* as it likely depends on longer tracks of single-stranded DNA^12-14^. Such tracks are typically available during repair of double-strand breaks and the majority of *kataegis* has been observed within 10kb of detected breakpoints^11^.

Amplification of known cancer genes due to double-strand breaks and complex rearrangements is known to drive tumorigenesis in many cancer types^33^. Recent studies have elucidated high copy number states of circular extrachromosomal DNA (ecDNA), which often harbor known cancer genes and have been found in most human cancers^33-36^. The circular nature of ecDNAs and their rapid replication patterns mimic double stranded DNA viral pathogens indicating a potential substrate for APOBEC3 mutagenesis, which may ultimately contribute to the subclonal diversification of tumors harboring ecDNA through accelerated diversification of the extrachromosomal oncoproteins.

Here we provide a comprehensive examination of clustered substitutions and clustered indels across 2,583 cancer genomes spanning 30 different tumor types. Our results elucidate a multitude of mutational processes giving rise to clustered mutations and reveal recurrent APOBEC3 mutagenesis fueling the evolution of ecDNA.

## RESULTS

### The landscape of clustered mutations in human cancer

To identify clustered mutations, an intra-mutational distance (IMD) cutoff was derived for each cancer sample where mutations below the cutoff were unlikely to occur by chance (q-value<0.01). A statistical approach utilizing the IMD cutoff, variant allele frequencies (VAFs), and corrections for local sequence context was applied to each examined specimen (**Methods**; **Extended Data Fig. 1*a***). Clustered mutations with consistent VAFs were subclassified into four categories (**Extended Data Fig. 1*b***). *Doublet-base substitutions* (DBSs) and *multi-base substitutions* (MBSs) were characterized, respectively, as 2 or ≥3 adjacent mutations (IMD=1). Multiple substitutions each with IMD above 1bp and below the sample-dependent cutoff were characterized as either *omikli* (2-3 substitutions) or *kataegis* (≥4 substitutions; **Supplementary Note 1**). Clustered substitutions with inconsistent VAFs were classified as *other*. While clustered indels were not subclassified into different categories (**Extended Data Fig. 1*b***), most events resembled diffuse hypermutation with 92.3% of events having only two indels and 97.9% having less than 4 indels (**Extended Data Fig. 1*c***).

Examining the 2,583 whole-genome sequenced cancers from the Pan-Cancer Analysis of Whole Genomes (PCAWG) project revealed a total of 1,686,013 clustered single-base substitutions and 21,368 clustered indels (**Fig. 1*a***). DBSs, MBSs, *omikli*, and *kataegis* comprise 45.7%, 0.7%, 37.2%, and 7.0% of clustered substitutions across all samples, respectively, with their distributions varying greatly within and across cancer types. For example, melanoma had the highest clustered substitution burden with ultraviolet light associated doublets (*viz*., CC>TT) accounting for 74.2% of clustered mutations; however, clustered substitutions contributed only 5.3% of all substitutions in melanoma (**Fig. 1*a***). In contrast, 11.5% of all substitutions in bone leiomyosarcomas were clustered with *omikli* and *kataegis* constituting 43.8% and 46.7% of these mutations, respectively (**Fig. 1*a***). Clustered indels exhibited similarly diverse patterns within and across cancer types (**Fig. 1*b***). For example, the highest mutational burden of clustered indels was observed in lung and ovarian cancers. Clustered indels in lung cancer accounted for only 2.6% of all indels and were characterized by 1bp deletions. In contrast, clustered long indels at microhomologies were commonly found in ovarian and breast cancers and contributed >10% of all indels in many samples (**Fig. 1*b***). Correlations between the total number of mutations and the number of clustered mutations were observed for DBSs and *omikli* but not for MBSs, *kataegis*, or indels (**Extended Data Fig. 1*d***). In most cancers, DBSs and *omikli* had VAFs consistent with the ones of non-clustered mutations while MBSs and *kataegis* tended to have lower VAFs (**Extended Data Fig. 1*e***). Distinct *kataegic* events contained 4 to 44 mutations with 81% of events being strand-coordinated, indicative of damage or enzymatic changes occurring on a single strand of DNA.

**Figure 1:**
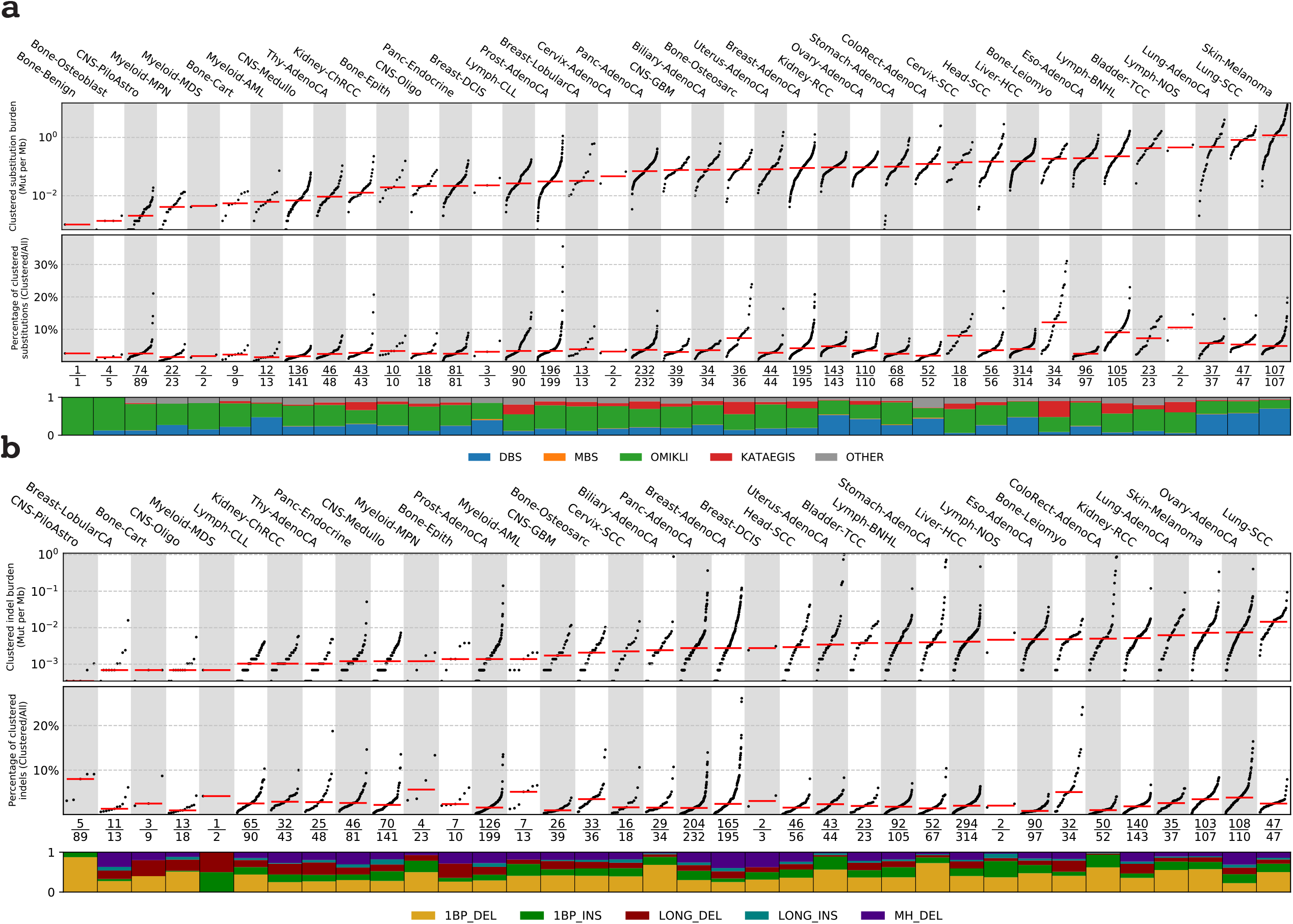
The landscape of clustered mutations across human cancer. ***a)*** Pan-cancer distribution of clustered substitutions subclassified into doublet-base substitutions, multi-base substitutions, *omikli, kataegis*, and *other* clustered mutations. *Top:* Each black dot represents a single cancer genome. Red bars reflect the median clustered tumor mutational burden (TMB) for cancer types. *Middle:* The clustered TMB normalized to the genome wide TMB reflecting the contribution of clustered mutations to the overall TMB of a given sample. Red bars reflect the median contribution for cancer types. *Bottom:* The proportion of each subclass of clustered events for a given cancer type with the total number of samples having at least a single clustered event over the total number of samples within a given cancer cohort. ***b)*** Pan-cancer distribution of clustered small insertions and deletions. *Top* and *Middle* panels have the same information as *a). Bottom:* The proportion of each cluster type of indel for a given cancer type with the total number of samples having at least a single clustered indel over the total number of samples within a given cancer cohort.

### Signatures of mutational processes giving rise to clustered mutations

To decipher the processes imprinting clustered mutations, mutational signature analysis was performed for each category of clustered events elucidating 12 DBS, 5 MBS, 17 *omikli*, 9 *kataegic*, and 6 clustered indel signatures (**Figs. 2*a***; **Supplementary Tables 1-5**). While DBS signatures have been previously described^1^, prior analysis combined DBSs and MBSs into a single class^1^. Separating these events into individual classes revealed that a multitude of mutational processes can give rise to DBSs while the majority of MBSs are attributable to signatures associated with tobacco smoking (SBS4) or ultraviolet light (SBS7). Additional MBS signatures were found within a small subset of cancers of the colon, esophagus, and head and neck (**Supplementary Note 1**).

**Figure 2:**
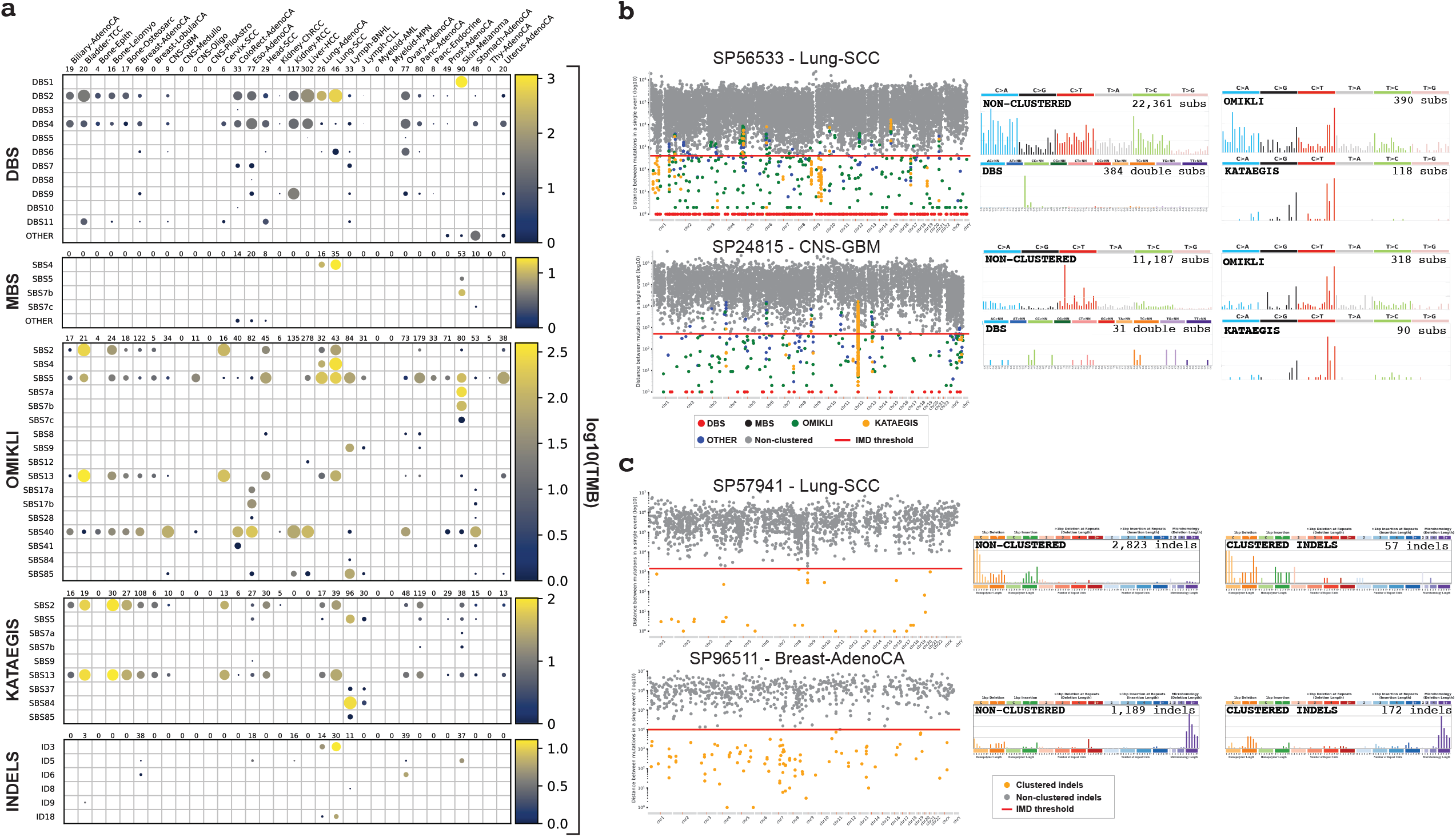
Mutational processes underlying clustered events. ***a*)** Each circle represents the activity of a signature for a given cancer type, where the radius of the circle determines the proportion of samples with greater than a given number of mutations specific to each subclass, while the color reflects the median number of mutations per cancer type. ***b*)** Two samples depicting the intra-mutational distance (IMD) distributions of substitutions across genomic coordinates, where each dot represents the minimum distance to adjacent mutations for a selected mutation colored based upon the corresponding subclassification of event (rainfall plot; *left*). The red lines depict the sample-dependent IMD threshold for each sample. Specific clustered mutations may be above this threshold based upon corrections for regional mutation density. The mutational spectra for the different catalogs of clustered and non-clustered substitutions for each sample (*right*; MBS are not shown). ***c*)** Two samples illustrating the IMD distributions of indels across the given genomes, with the IMD indel thresholds shown in red (*left*). The non-clustered and clustered indel catalogs for each sample (*right*).

In cancer genomes, *omikli* were previously attributed to APOBEC3 mutagenesis^15^ with some indirect evidence from experimental models^24,37-39^. Our analysis of sequencing data^40^ from the clonally expanded breast cancer cell line BT474 with active APOBEC3 mutagenesis experimentally confirmed the existence of APOBEC3 *omikli* events (cosine similarity: 0.99; **Extended Data Fig. 2*a***). Only 16.2% of *omikli* events across the 2,583 cancer genomes matched the APOBEC3 mutational pattern indicating that a plethora of other mutational processes can give rise to diffuse clustered hypermutation. In addition to APOBEC3-associated events, our mutational signature analysis revealed *omikli* due to tobacco smoking (SBS4), clock-like mutational processes (SBS5), ultraviolet light (SBS7), both direct and indirect mutations from AID (SBS9 and SBS85), and multiple mutational signatures with unknown etiology in different cancer types (SBS8, SBS12, SBS17a/b, SBS28, SBS40, and SBS41; **Fig. 2*a***). Cell lines previously exposed to benzo[*a*]pyrene^41^ and ultraviolet light^42^ confirmed the generation of *omikli* events due to these two environmental exposures (cosine similarities: 0.86 and 0.84, respectively; **Extended Data Fig. 2*a***).

From the 9 *kataegic* signatures, 4 have been reported previously including 2 associated with the activity of APOBEC3 deaminases (SBS2 and SBS13) and 2 associated with both canonical and non-canonical activities of AID (SBS84 and SBS85; **Fig. 2*a***). SBS5 (clock-like mutagenesis) accounted for 15.0% of observed *kataegis* with majority of these events occurring in the vicinity of AID *kataegis* within CLL or B-cell lymphomas. The remaining 4 *kataegic* signatures accounted for only 8.9% of *kataegic* mutations and included: SBS7a/b (ultraviolet light), SBS9 (indirect mutations from AID), and SBS37 (unknown etiology). Most *kataegic* signatures were strand-coordinated including SBS2, SBS5, SBS7b, SBS13, SBS84, and SBS85 (**Extended Data Fig. 2*b***). Notably, we observed a strand-coordinated APOBEC3 activity within a subset of glioblastomas, which had been missed by prior studies (**Fig. 2*b***). Some samples exhibited consistent while others exhibited distinct signatures of clustered and non-clustered mutagenesis (**Fig. 2*b***). For example, in SP56533 (lung squamous cell carcinoma), most non-clustered and *omikli* substitutions were caused by tobacco signature SBS4, while *kataegic* events were generated by the APOBEC3 signatures (**Fig. 2*b***). In contrast, the pattern of non-clustered substitutions in SP24815 (glioblastoma) was due to clock-like signatures SBS1 and SBS5 while the predominant *omikli* and *kataegic* events were attributable to APOBEC3 (**Fig. 2*b***).

The remaining *other* clustered substitutions exhibited inconsistent VAFs likely representing mutations at highly mutable genomic regions or the effects of co-occurring large mutational events such as copy number alterations (**Extended Data Fig. 2*d***; **Supplementary Table 6**).

Different cancers revealed distinct tendencies of clustered indel mutagenesis (**Fig. 2*a***). For instance, clustered indels attributed to ID3 (tobacco smoking; characterized by 1bp deletions) were found predominately in lung cancers and significantly elevated in smokers compared to non-smokers (p-value: 0.0014; **Fig. 2*c***; **Extended Data Fig. 2*c***). Importantly, clustered indels due to signatures ID6 and ID8, both attributed to homologous recombination deficiency and characterized by long indels at microhomologies, were found in breast and ovarian cancers and were highly elevated in cancers with known deficiencies in homologous recombination genes (p-value: 4.9 × 10^−11^; **Fig. 2*c***; **Extended Data Fig. 2*c***).

### The panorama of clustered driver mutations in human cancer

The PCAWG project elucidated a constellation of substitutions and indels with a high confidence of being mutations that drive cancer development^11^. Our current analysis reveals statistically significant enrichments of clustered substitutions and clustered indels amongst these driver mutations. Specifically, whereas only 3.7% of all substitutions and 0.9% of all indels are clustered events, they contribute 8.4% and 6.9% of substitution and indel drivers, respectively (q-values<1e-5; Fisher’s exact tests; **Figure 3*a*&*b***). *Omikli* accounted for 50.5% of all clustered substitution drivers, while DBSs, *kataegis*, and *other* clustered events each contributed between 14% and 18% (**Figure 3*c***). The prevalence of clustered driver substitutions varied greatly between genes and across different cancers (**Figure 3*c***; **Extended Data Fig. 3*a***). In some cancer genes, only a small percentage of driver events are due to clustered substitutions; examples include *TP53* (4.5% clustered driver substitutions; 35 clustered substitutions out of a total of 772 driver substitutions), *KRAS* (3.7%; 10/270), and *PIK3CA* (2.2%; 4/178). Nevertheless, in other genes, most detected substitution drivers were clustered events; examples include: *BTG1* (73.1%; 19/26), *SGK1* (66.6%; 6/9), *EBF1* (60.0%; 6/10), and *NOTCH2* (38.5%; 5/13). Importantly, the contribution from each class of clustered events varied across the driver substitutions in different genes (**Figure 3*c***). For instance, ultraviolet light associated DBSs comprise 93% of clustered *BRAF* driver events, *omikli* contribute 63% of clustered *BTG1* driver events, and *kataegis* accounted for 100% of clustered *NOTCH2* driver substitutions (**Figure 3*c***). Similar behavior was observed for clustered indel drivers with 48.7% of all clustered indel drivers being single-base pair indels (**Figure 3*b***). In some cancer genes, clustered indel drivers were rare events (*e*.*g*., 2.4% of indel drivers in *TP53* were clustered indels) whereas in others they were commonly found (*e*.*g*., 76.6% in *ALB*; **Figure 3*d***). The compendium of clustered driver events was induced by the activity of multiple different mutational processes including exposure to ultraviolet light, tobacco smoke, platinum chemotherapy, and AID/APOBEC3 activity; amongst others (**Extended Data Fig. 3*b***).

**Figure 3:**
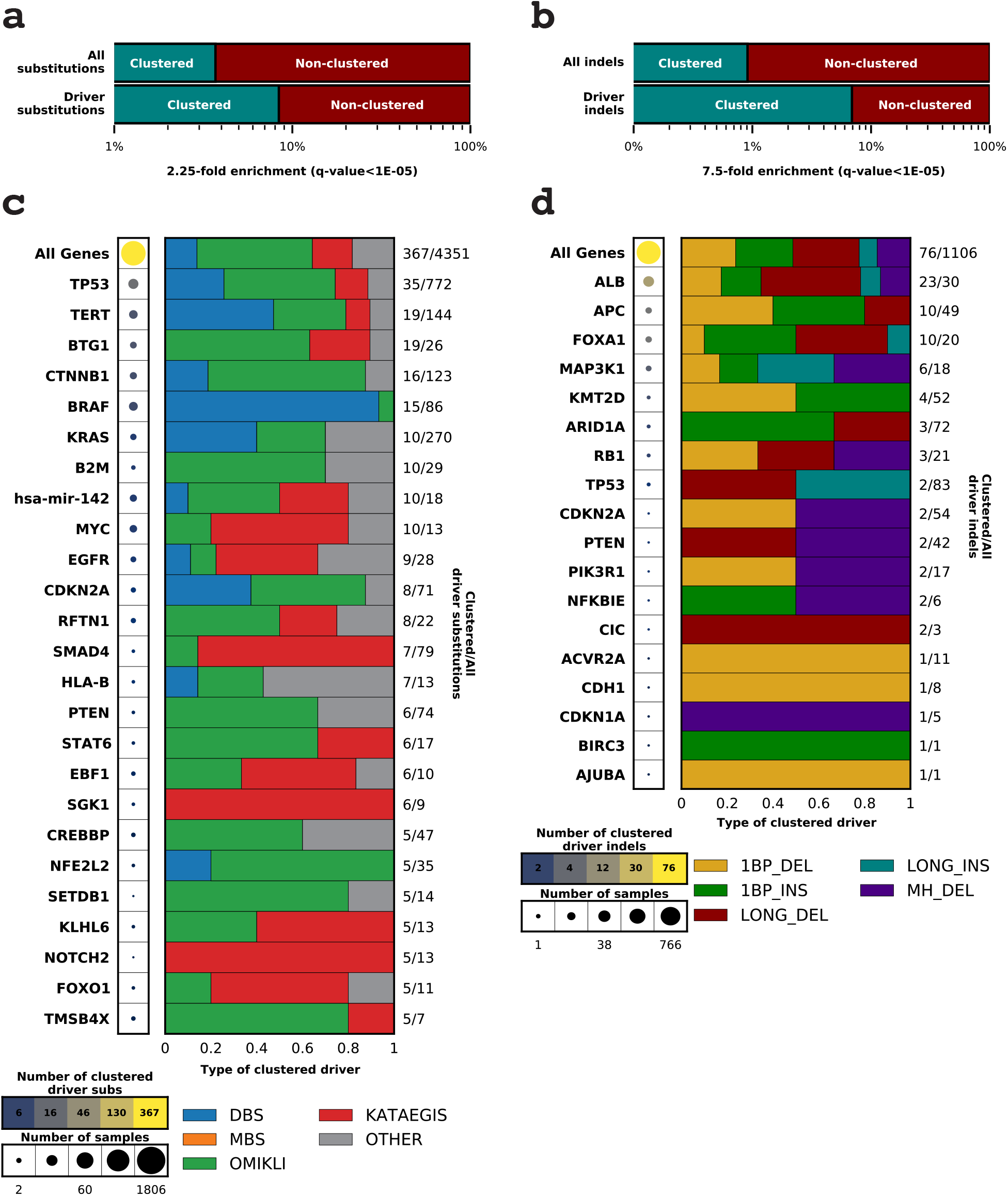
Panorama of clustered driver mutations in human cancer. Percentage of clustered mutations (*top*) compared to the percentage of clustered driver events (*bottom*) for substitutions ***a*)** and indels ***b*)**. P-values were calculated using a Fisher’s exact test and corrected for multiple hypothesis testing using Benjamini-Hochberg false discovery rate procedure. ***c*)** The frequency of clustered driver events across known cancer genes. The radius of the circle is proportional to the number of samples with a clustered driver mutation within a gene, while the color reflects the clustered mutational burden. All clustered driver events are classified into one of the five clustered classes, with the number of clustered driver substitutions and the total number of driver substitutions shown on the right. ***d*)** Clustered indels drivers are displayed in a format similar to ***c)***.

### *Kataegic* events associate with a variety of focal amplifications and structural variations

In each sample, *kataegic* mutations were separated into distinct events based on consistent VAFs across adjacent mutations as well as IMD distances greater than the sample-dependent IMD threshold (**Methods**). Our analysis revealed that 36.2% of all *kataegic* events occurred within 10kb of a structural breakpoint but not on detected focal amplifications (**Fig. 4*a***). Additionally, 21.8% of all *kataegic* events occurred either on a detected focal amplification or within 10kb of a focal amplification’s structural breakpoint: 9.6% on circular extrachromosomal DNA (ecDNA), 6.3% on linear rearrangements, 3.3% within heavily rearranged events, and 2.6% associated with BFBs (**Fig. 4*a***). Lastly, 42.0% of *kataegic* events were neither within 10kb of a structural breakpoint nor on a detected focal amplification. Modelling the distribution of the distances between *kataegic* events and the nearest structural variations revealed a multi-modal distribution with three components (**Fig. 4*b***): *(i) kataegis* within 10kb of a breakpoint; *(ii) kataegis* within ∼10Mb of a breakpoint; *(iii) kataegis* within >1.5Mb from a nearby breakpoint (similar distribution to non-clustered mutations). Importantly, ecDNA-associated *kataegis* had ∼750kb average distance from the nearest breakpoint with only 0.35% of *kataegic* events occurring both on ecDNA and within 10kb of a breakpoint (**Fig. 4*b***). These results indicate that ecDNA-associated *kataegic* events are not likely to have occurred due to structural rearrangements during the formation of ecDNA. In most cancer types, DBSs, MBSs, *omikli*, and other cluster events were not found in the vicinity of structural variations (**Extended Data Fig. 4*a&b***).

**Figure 4:**
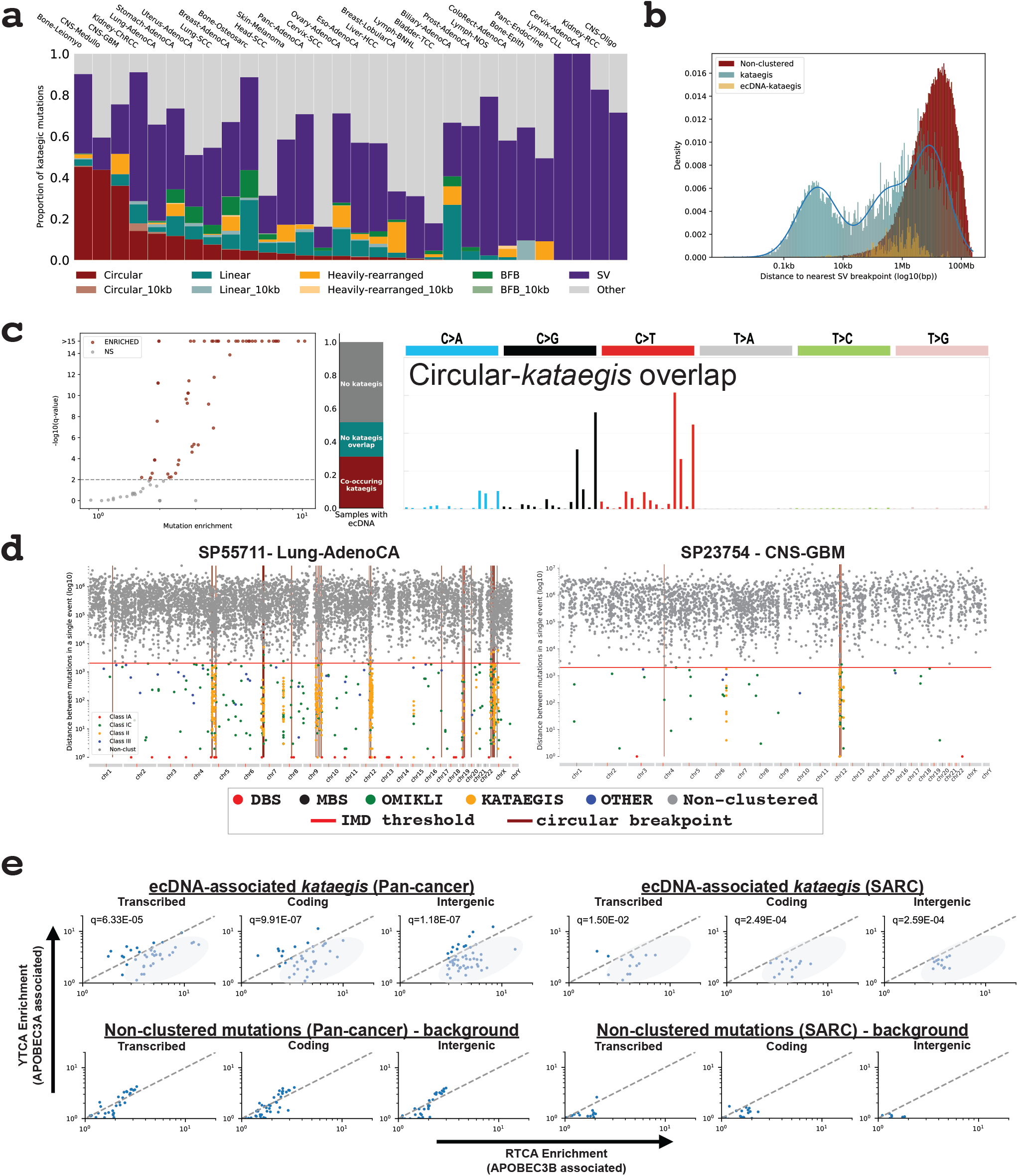
*Kataegic* events co-locate with most forms of structural variations. ***a*)** The proportion of all *kataegic* events for each cancer type overlapping different forms of focal amplification or structural variations. ***b*)** The distance to the nearest breakpoint for all *kataegic* mutations (teal), *kataegic* mutations (gold) found on ecDNA, and non-clustered mutations (red). The distribution for all *kataegis* distances were modeled using a Gaussian mixture comprised of three components. ***c*)** *Left:* Volcano plot depicting a subset of samples that are statistically enriched for *kataegis* mutations within ecDNA regions (red; q-values determined using a Z-test). *Middle:* The proportion of samples with ecDNA that co-occur with *kataegis*, do not co-occur with *kataegis*, or do not have any detected *kataegic* activity. *Right*: The mutational spectrum of all *kataegic* somatic mutations co-occurring *with* ecDNA. ***d*)** Rainfall plots illustrating the intra-mutational distance (IMD) distribution for a given sample with the genomic locations of ecDNA breakpoints shown in maroon. ***e*)** YT<u>C</u>A versus RT<u>C</u>A enrichments per sample within ecDNA *kataegis*, where YT<u>C</u>A and RT<u>C</u>A enrichment is suggestive of higher APOBEC3A or APOBEC3B activity, respectively. Genic mutations were divided into transcribed (template strand) and coding mutations. The RT<u>C</u>A/YT<u>C</u>A fold enrichments were compared to the fold enrichments of non-clustered mutations (p-values calculated using a Mann-Whitney *U*-tests and corrected for multiple hypothesis testing using the Benjamini-Hochberg false discovery rate procedure).

### Multiple distinct *kataegic* events suggest recurrent mutagenesis of ecDNA

While only 9.6% of *kataegic* events occur within ecDNA regions, >30% of ecDNAs had one or more associated *kataegic* events (**Fig. 4*c***). The somatic mutations within these ecDNA regions were dominated by the APOBEC3 mutational patterns, which are characterized by strand-coordinated C>G and C>T mutations at Tp<u>C</u>pW context and attributed to signatures SBS2 and SBS13 (**Fig 4*c*&*d*; Extended Data Fig. 4*c***). These APOBEC3-associated events contributed 97.8% of all *kataegic* events within ecDNA, while the remaining mutations were attributed to clock-like signature SBS5 (1.2%) and other mutational signatures (1.0%). Further, ecDNA-associated *kataegic* events exhibited a high enrichment of C>T and C>G mutations at RT<u>C</u>A compared to YT<u>C</u>A contexts^12^ indicating that APOBEC3B plays an important role in the mutagenesis of circular DNA bodies (**Fig. 4*e***). Similar levels of enrichment for RT<u>C</u>A contexts are also observed in both non-ecDNA *kataegis* and non-SV associated *kataegis* suggesting that APOBEC3B generally gives rise to many of the strand-coordinated *kataegic* events (**Extended Data Fig. 4*d***).

More recurrent APOBEC3 *kataegis* was observed across circular ecDNA regions compared to other forms of structural variations (**Fig. 5*a***). An average of 2.5 distinct *kataegic* events were observed within ecDNA regions (range: 0 to 64 *kataegic* events; 0 to 505 mutations). Recurrent APOBEC3 mutagenesis of ecDNA was widespread across distinct cancer types. For instance, glioblastomas and sarcomas exhibited an average of 5 and 86 *kataegic* APOBEC3-like mutations overlapping ecDNA, respectively. The average VAF of ecDNA-associated *kataegis* was significantly lower than both non-ecDNA associated *kataegis* and all other clustered events (q-values<1e-5; **Fig. 5*b***). Interestingly, a subset of ecDNA-associated *kataegis* exhibited VAFs above 0.80 likely reflecting early mutagenesis of genomic regions that have subsequently amplified as ecDNA. Further, *kataegic* events with high VAFs occurred more commonly on ecDNA harboring known cancer genes suggesting a mechanism of positive selection (**Fig. 5*b***). We estimate that ∼7.2% of ecDNA-associated *kataegis* occurred early in the evolution of a given ecDNA population within a tumor (VAF>0.80), while the majority of ecDNA-associated *kataegic* events (∼82.5%; VAF<0.5) have likely occurred after clonal amplification by recurrent APOBEC3 *kataegic* mutagenesis.

**Figure 5:**
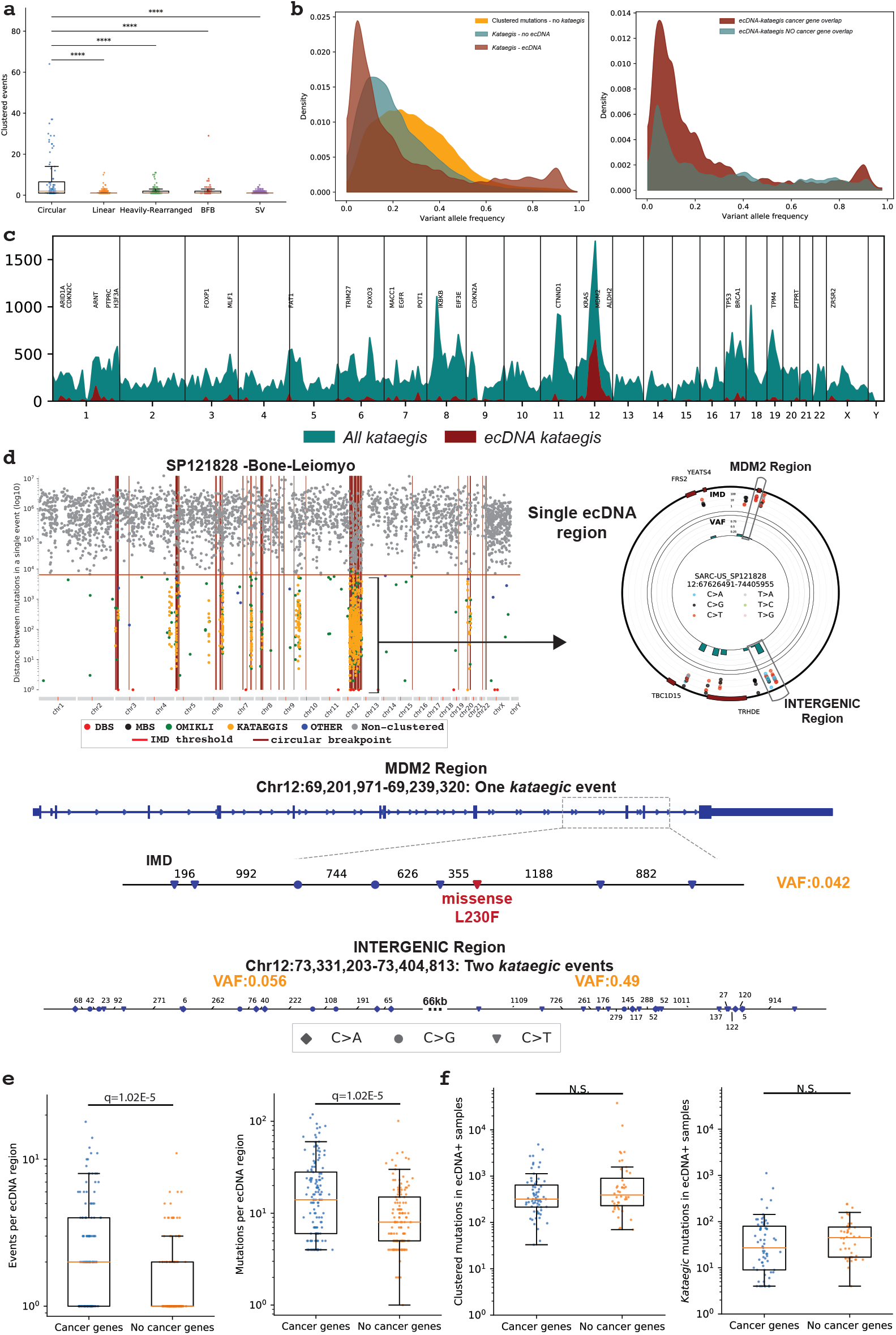
Recurrent APOBEC3 hypermutation of ecDNA. ***a*)** The number of clustered events overlapping a single structural variation event, where each dot represents a single structural variation. A 10kb window was used to determine the co-occurrence of *kataegis* with simple structural variation breakpoints (SV; purple). ***b*)** *Left:* The normalized distributions of the variant allele frequencies for all clustered mutations excluding *kataegis* (orange), all non-ecDNA *kataegis* (teal), and ecDNA *kataegis. Right:* The normalized distributions of the variant allele frequencies for ecDNA *kataegis* harboring known cancer genes and for ecDNA *kataegis* that do not harbor known cancer genes. ***c*)** The frequency of recurrence for all *kataegis* (teal) and ecDNA *kataegis* (red) across the genome calculated using a sliding window of 10Mb. Known cancer genes within proximity to frequency peaks of ecDNA *kataegis* are displayed. ***d*)** A single sarcoma genome depicting the overlap of *kataegis* with ecDNA regions displayed as a rainfall (*top left)* with a single zoomed in ecDNA represented using a circos plot (*top right*). The outer track of the circos plot represents the reference genome of the ecDNA with proximal known cancer driver genes. The middle track reflects a circular rainfall plot where each dot represents the intra-mutational distance (IMD) around a single mutation colored based on the substitution change. The innermost track shows the average variant allele frequency (VAF) for each *kataegic* event. *Bottom*: Two smaller regions of the selected ecDNA including a single *kataegic* event within *MDM2* region resulting in a missense mutation, and two *kataegic* events within an intergenic region with the average VAF per event (orange). ***e*)** The number of *kataegic* events and total *kataegic* mutations per ecDNA region that harbor known cancer genes and that do not harbor known cancer genes (*left* and *right*; respectively). ***f*)** The total number of clustered mutations and *kataegic* mutations found in samples with ecDNAs containing cancer genes compared to samples with ecDNAs without any known cancer genes (*left* and *right*; respectively). P-values were calculated using a Mann-Whitney U test and corrected using the Benjamini-Hochberg false discovery rate procedure.

To investigate the oncogenic effect of ecDNA-associated *kataegis*, we examined regions of recurrence across all cancer samples. The ecDNA-associated *kataegis* showed an increased mutational burden within or near known cancer-associated genes including *TP53, CDK4*, and *MDM2*, amongst others (**Fig. 5*c***). These events were observed across many cancers including glioblastomas, sarcomas, head and neck squamous cell carcinomas, and lung adenocarcinomas suggesting a recurrent ecDNA-associated *kataegis* (**Fig. 5*d***; **Extended Data Fig. 5*a***). For example, in a sarcoma sample (SP121828), 10 distinct *kataegic* events overlapped a single ecDNA region with recurrent APOBEC3 activity in proximity to *MDM2* resulting in a missense L230F mutation (**Fig. 5*d***). The same ecDNA region harbored additional *kataegic* events occurring within intergenic regions that have distinguishable VAF distributions implicating recurrent mutagenesis (**Fig. 5*d***). Similarly, two distinct *kataegic* events occur on an ecDNA harboring *EGFR* resulting in a missense mutation D191N within a head and neck cancer (**Extended Data Fig. 5*a***). Importantly, ecDNA regions with known cancer-associated genes had a significantly higher number of *kataegic* events and a significantly higher mutational burden compared to ecDNA regions without any known cancer-associated genes (q-values<1e-5; **Fig. 5*e***). Further, we observed a higher co-occurrence of *kataegis* and ecDNA with known cancer-associated genes, which were mutated 2.5 times more than ecDNA without cancer-associated genes (p-value=1.2e-5; Fisher’s exact test). Overall, 41% of *kataegic*-ecDNA events were found within the footprints of known cancer driver genes (p-value<1e-5). These enrichments cannot be accounted either by an increase in the overall mutations or by an increase in the overall clustered mutations in these samples (**Fig. 5*f***). To understand the functional effect of ecDNA-associated *kataegis*, we annotated the predicted consequence of each mutation. In total, 2,247 *kataegic* mutations overlapped putative cancer-associated genes, of which 4.3% occur within coding regions (**Extended Data Fig. 5*b***). Specifically, 63 resulted in missense mutations, 29 resulted in synonymous mutations, 4 introduced premature stop codons, and 1 removed a stop codon (**Supplementary Table 7**). These downstream consequences of APOBEC3 mutagenesis suggest a contribution to the oncogenic evolution of specific ecDNA populations.

## DISCUSSION

Clustered mutagenesis in cancer can occur through multiple different mutational processes, with AID/APOBEC3 deaminases playing the most prominent role. In addition to enzymatic deamination, other endogenous and exogenous sources imprint many of the observed clustered indels and substitutions. Importantly, a multitude of mutational processes can give rise to *omikli* events including tobacco carcinogens and exposure to ultraviolet light. Cluster substitutions and indels are 2.3-fold and 7.7-fold enriched in driver events, respectively, implicating them in cancer development and cancer evolution. Some clustered mutational signatures are associated with known cancer risk factors as well as the activity or failure of DNA repair processes.

The large proportion of *kataegic* events occur within 10kb of detected breakpoints with a mutational pattern suggesting the activity of APOBEC3. Multiple distinct *kataegic* events, independent of detected breakpoints, were observed on ecDNA implicating recurrent APOBEC3 mutagenesis. The circular topology of ecDNAs and their rapid replication patterns are reminiscent of the structure and behavior of the circular genomes of several double stranded DNA based pathogens including herpesviruses, papillomaviruses, and polyomaviruses^33-36^. Importantly, prior pan-virome studies have shown that these double stranded DNA viral genomes often manifest mutations from APOBEC3A/B enzymes^43-45^. As such, recurrent APOBEC3 mutagenesis on ecDNA is likely representative of an anti-viral response where the ecDNA viral-like structure is treated as an infectious agent and attacked by APOBEC3A/B enzymes. ecDNAs harbor a plethora of cancer-associated genes and are responsible for many gene amplification events that can accelerate tumor evolution. Repeated mutagenic attacks of these ecDNA reveals functional effects within known oncogenes implicating additional modes of oncogenesis that may ultimately contribute to subclonal tumor evolution and subsequent evasion to therapy. Further investigation is required to fully understand the clinical implications of APOBEC3 mutagenesis of ecDNA.

## ONLINE METHODS

### Data Sources

Somatic variant calls of single-base substitutions, small insertions and deletions, and structural variations were downloaded for the 2,583 white-listed whole-genome sequenced samples from PCAWG along with the corresponding list of consensus driver events^11^. Epidemiological and clinical features for all available samples were downloaded from the official PCAWG release (https://dcc.icgc.org/releases/PCAWG). The subclassification of focal amplifications comprised of circular extrachromosomal DNA (ecDNA), linear amplifications, breakage-fusion-bridge cycles (BFBs), and heavily rearranged events, and their corresponding genomic locations were obtained for a subset of samples (*n* = 1,291) using AmpliconArchitect as reported^35^.

Experimental models used to validate clustered events were derived from previous studies using primary Hupki mouse embryonic fibroblasts (MEFs) exposed to ultraviolet light^42^, human induced pluripotent stem cells (iPSC) exposed to benzo[*a*]pyrene^41^, and clonally expanded BT-474 human breast cancer cell line with episodically active APOBEC3^40^.

### Detection and classification of clustered mutational events

SigProfilerSimulator (v1.0.2) was used to derive an intra-mutational distance (IMD) cutoff^46^ that is unlikely to occur by chance based upon the tumor mutational burden and the mutational patterns for a given sample. Specifically, each tumor sample was simulated while maintaining the sample’s mutational burden on each chromosome, the +/-2bp sequence context for each mutation, and the transcriptional strand bias ratios across all mutations. All mutations in each sample were simulated 100 times and the IMD cutoff was calculated such that 90% of the mutations below this cutoff could not appear by chance (q-value<0.01). For example, in a sample with an IMD threshold of 500bp, one may observe 1,000 mutations within this threshold with no more than 100 mutations expected based on the simulated data (q-value<0.01). P-values were calculated using z-tests by comparing the number of real mutations and the distribution of simulated mutations that occur below the same IMD threshold. A maximum cutoff of 10kb was used for all IMD thresholds. By generating a background distribution that reflects the random distribution of events used to reduce the false positive rate, this model also considers regional heterogeneities of mutation rates and variances in clonality by correcting for mutation-rich regions within 1Mb windows and ensuring that subsequent mutations likely occurred as single events using a maximum cutoff of 0.10 for differences in the variant allele frequencies (VAFs). The regional IMD cutoff was determined using a sliding window approach that calculated the fold enrichment between the real and simulated mutation densities within 1Mb windows across the genome. The IMD cutoffs were further increased, for regions that had higher than 9-fold enrichments of clustered mutations and where >90% of the clustered mutations were found within the original data, to capture additional clustered events while maintaining the original criteria (<10% of the mutations below this cutoff appear by chance; q-value<0.01).

Subsequently, clustered substitutions were classified into a single type of clustered event. All clustered mutations with consistent VAFs were classified into one of four categories (**Extended Data Fig. 1*a***). Two adjacent mutations with an IMD of 1 were classified as *doublet-base substitutions*. Three or more adjacent mutations each with an IMD of 1 were classified as *multi-base substitutions*. Two or three mutations with IMDs less than the sample-dependent threshold and with at least a single IMD greater than 1 were classified as *omikli*. Four or more mutations with IMDs less than the sample-dependent threshold and with at least a single IMD greater than 1 were classified as *kataegis*. A cutoff of four mutations for *kataegis* was chosen by fitting a Poisson mixture model to the number of mutations involved in a single event across all extended clustered events excluding DBSs and MBSs (**Supplementary Note 1**). This model comprised two distributions with C1=2.08 and C2=4.37 representing *omikli* and *kataegis*, respectively. A cutoff of four mutations was used for *kataegis* based upon >95% contribution from the *kataegis*-associated distribution with events of four or more mutations. Note that there is certain ambiguity for events with 2 or 3 mutations. While the majority of these events are *omikli*, some of these events are likely short *kataegic* events (**Supplementary Note 1**). All remaining clustered mutations with inconsistent VAFs were classified as *other*. Clustered indels were not classified into different classes.

### Analysis of clustered mutational patterns and clustered mutational signatures

The clustered mutational catalogues of the examined samples were summarized in SBS288 and ID83 matrices using SigProfilerMatrixGenerator^47^ for each tissue type and each category of clustered events. For example, six matrices were constructed for clustered mutations found in Breast-AdenoCA: one matrix for DBSs, one matrix for MBSs, one matrix for *omikli*, one matrix for *kataegis*, one matrix for other clusters substitutions, and one matrix for clustered indels. The SBS288 classification considers the 5’ and 3’ bases immediately flanking each single-base substitution (referred to using the pyrimidine base in the Watson-Crick base pair) resulting in 96 individual mutation channels. Further, this classification considers the strand orientation for mutations that occur within genic regions resulting in three possible categories; *(i)* transcribed; pyrimidine base occurs on the template strand; *(ii)* untranscribed; pyrimidine base occurs on the coding strand; or *(iii)* non-transcribed; pyrimidine base occurs in an intergenic region. Note that mutations in genic regions that are bi-directionally transcribed were evenly split amongst the coding and template strand channels. Combined, this results in a classification consisting of 288 mutation channels, which were used as input for *de novo* signature extraction of clustered substitutions. The ID83 mutational classification has been previously described^47^.

Mutational signatures were extracted from the generated matrices using SigProfilerExtractor (v1.1.0), a Python based tool that uses nonnegative matrix factorization to decipher both the number of operative processes within a given cohort and the relative activities of each process within each sample^48^. The algorithm was initialized using random initialization and by applying multiplicative updates using the Kullback-Leibler divergence with 500 replicates. Each *de novo* extracted mutational signature was subsequently decomposed into the COSMIC (v3) set of signatures (https://cancer.sanger.ac.uk/signatures/) requiring a minimum cosine similarity of 0.80 for all reconstructed signatures. All *de novo* extractions and subsequent decomposition were visually inspected and, as previously done^1^, manual corrections were performed for 2.2% of extractions (4 out of 180 extractions) where the total number of operative signatures was adjusted ±1. Consistent with prior visualizations^1^, decomposed signature activity plots required that each cancer type have more than 2 samples and used mutation thresholds for each clustered category; 25 mutations per sample were required for doublet-base substitutions, *omikli* events, and other clustered mutations; 15 mutations per sample were required for multi-base substitutions and *kataegic* events; 10 mutations were required per sample for clustered indels.

### Experimental validation of clustered signatures

A subset of clustered mutational signatures was validated using previously sequenced *in vitro* cell line models. As done for PCAWG samples, we generated a background model using SigProfilerSimulator^46^ to calculate the clustered IMD cutoff for each sample and partitioned each substitution into the appropriate category of clustered events. Mutational spectra were generated for each subclass within each sample using SigProfilerMatrixGenerator^47^ and were compared against the *de novo* signatures extracted from human cancer. The cosine similarity between the *in vitro* mutational spectra and *de novo* observed clustered signatures was calculated to assess the degree of similarity. Note that cosine similarities above 0.81 reflect p-values below 0.01 (Ref. ^46^).

### Associations between clustered mutational signatures and cancer risk factors

Homologous recombination deficiency (HRD) was defined for breast cancers using the status of *BRCA1, BRCA2, RAD51C*, and *PALB2*^49^. Samples with a germline, somatic, or epigenetic alteration in one of these genes were considered HR-deficient, while samples without any known alterations in these genes were considered HR-proficient. The number of clustered indels were compared between HR-deficient and HR-proficient samples. The smoking status of lung cancers was determined using the clinical annotation from TCGA (https://portal.gdc.cancer.gov/repository). The number of clustered indels associated with tobacco smoking (ID6) were compared between samples annotated as lifelong non-smokers and samples annotated as current and reformed smokers. The status of alcohol consumption was determined using the annotations from the official PCAWG release (https://dcc.icgc.org/releases/PCAWG). The total number of clustered indels were compared in samples annotated with no alcohol consumption and those annotated as daily and weekly drinkers.

### Modelling the distributions of distances between structural variations and clustered events

The distance to the nearest structural variation breakpoint was calculated for each mutation in each subclass using the minimum distance to the nearest adjacent upstream or downstream breakpoint. Each distribution was modeled using a Gaussian mixture with an automatic selection criterion for the number of components ranging between one and five components using the minimum Bayesian information criteria (BIC) across all iterations. Modelling of *kataegic* events resulted in an optimal fit of three components, which was used to separate *kataegic* substitutions into SV-associated and non-SV associated mutations for subsequent analysis. Doublet-base substitutions and multi-base substitutions were both modelled using a single Gaussian distribution relating to non-SV associated mutations, while *omikli* and other clustered mutations were modelled using a mixture of two components likely reflecting leakage of smaller *kataegic* events contributing to a weak SV-associated distribution.

### Enrichment analysis of APOBEC3A and APOBEC3B mutagenesis (RT<u>C</u>A *versus* YT<u>C</u>A)

The enrichment score of RT<u>C</u>A and YT<u>C</u>A penta-nucleotides quantifies the frequency for which each Tp<u>C</u>pA>Tp<u>K</u>pA mutation occurs at either an RT<u>C</u>A or YT<u>C</u>A context. To account for motif availability, this score is calculated using the +/-20bp sequence context around each mutation and normalized by the number of cytosine bases and C>N mutations within the set of 41-mers surrounding each mutation of interest^12^.

### Associations between circular ecDNA and *kataegis*

The collection of ecDNA ranges were intersected with the catalog of clustered mutations, which was used to determine the overlapped mutational burden for each subclass of clustered event and the mutational spectra of overlapping *kataegic* events. Enrichments of events were calculated using statistical background models generated using SigProfilerSimulator^46^ that shuffled the dominant mutation in each clustered event across the genome (*i*.*e*., the most frequent mutation type in a single event). Comparisons between ecDNA with and without cancer genes were performed using the set of cancer genes from the Cancer Gene Census (CGC)^50^. All statistical comparisons and p-values were calculated using a Mann-Whitney U test unless otherwise specified. For each set of tests, p-values were corrected for multiple hypothesis testing using the Benjamini-Hochberg false discovery rate procedure. The predicted effect of each overlapping variant was determined using ENSEMBL’s Variant Effect Predictor tool by reporting only the most severe consequence^51^.

## Data Availability

No data were generated specifically for this study. All data were and can be downloaded from the appropriate references.

## Code Availability

The SigProfiler compendium of tools are developed as Python packages and are freely available for installation through PyPI or directly through GitHub (https://github.com/AlexandrovLab/). For all tools, each package is fully functional, free, and open sourced distributed under the permissive 2-Clause BSD License and are accompanied by extensive documentation: *(i)* SigProfilerMatrixGenerator (https://osf.io/s93d5/wiki/home/); *(ii*) SigProfilerSimulator: (https://osf.io/usxjz/wiki/home/); *(iii*) SigProfilerExtractor: (https://osf.io/t6j7u/wiki/home/). Each tool also has an R wrapper available for installation through the GitHub repositories. The core computational pipelines used by the PCAWG Consortium for alignment, quality control and variant calling are available to the public at https://dockstore.org/search?search=pcawg under the GNU General Public License v.3.0, which allows for reuse and distribution.

## ACKNOWLEDGEMENTS

ENB and LBA were supported by Cancer Research UK Grand Challenge Award C98/A24032 as well as US National Institute of Health grants R01ES030993-01A1 and R01ES032547. LBA is an Abeloff V Scholar and he is also supported by an Alfred P. Sloan Research Fellowship. Research at UC San Diego was also supported by a Packard Fellowship for Science and Engineering to LBA. MP is supported by the European Molecular Biology Organization (EMBO) Long-Term Fellowship (ALTF 760-2019). VB is supported in part by a grant from the NIH (GM114362). Cancer research in the Harris lab is supported by NCI P01 CA234228. RSH is the Margaret Harvey Schering Land Grant Chair for Cancer Research, a Distinguished University McKnight Professor, and an Investigator of the Howard Hughes Medical Institute.

The funders had no roles in study design, data collection and analysis, decision to publish, or preparation of the manuscript.

## AUTHOR CONTRIBUTIONS

ENB and LBA designed the overall study. ENB performed all analyses with help from JL, MP, VB, PSM, RSH, and LBA. ENB and LBA wrote the manuscript with help and input from all other authors. All authors read and approved the final manuscript.

## EXTENDED DATA FIGURE LEGENDS

**Extended Data Figure 1: Identification and classification of clustered events. *a*)** Schematic depiction of the approach for separating clustered mutations from the complete mutational catalog for a given sample. ***b*)** The subclassification for all clustered substitutions and indel events. The expected intra-mutational distance (IMD) is calculated using steps 2 and 3 of panel *a*. ***c*)** The distribution of the number of indels present in a single clustered event. ***d*)** The correlations of the total tumor mutational burden (TMB) of each sample, the TMB within the exome, or the TMB for each class of clustered substitutions (*left*) and indels *(right*). ***e*)** The distribution of variant allele frequencies for all clustered substitution classes (*left*) with the average fold enrichment compared against the distribution of non-clustered mutations (*right*).

**Extended Data Figure 2: Experimental validation and epidemiological associations of clustered mutational processes. *a*)** Experimental validation of three *omikli* processes.

Specifically, APOBEC3-associated *omikli* were validated using a clonally expanded BT-474 breast cancer cell line (*top*), *omikli* resulting from exposure to benzo[*a*]pyrene were validated using iPSC cells (*middle*), and *omikli* resulting from exposure to ultraviolet light were validated using iPSC cells (*bottom*). ***b*)** Mutational processes of strand-coordinated *kataegic* events. ***c*)** Epidemiological associations comparing the ratio of clustered tumor mutational burden (TMB) to the total TMB for a given sample between drinkers and non-drinkers, smokers and non-smokers, and homologous-recombination deficient and homologous proficient samples. ***d*)** Mutational processes of clustered events with inconsistent variant allele frequency (VAFs) classified as *other* clustered substitutions.

**Extended Data Figure 3: Mutational processes of clustered driver events. *a*)** The percentage of clustered driver substitutions and indels within each cancer type. ***b*)** The proportional activity of mutational signatures contributing to clustered driver events within each subclass. Multi-base substitutions (MBSs) did not contribute to any reported driver events.

**Extended Data Figure 4: Clustered events and structural variations. *a*)** The proportion of all clustered events co-locating with structural variations across all cancer types (*left*) and across each cancer type (*right*). ***b*)** The distance to the nearest structural variation for each class of clustered mutations (teal), and non-clustered mutations (red). The distribution for each class of clustered events were modeled using a Gaussian mixture. DBSs and MBSs were modeled using a single distribution, while *omikli, other*, and indels were modeled using two components reflecting the minimal distribution of overlap with structural variations. ***c*)** The mutational signatures active in ecDNA clustered events. ***d*)** YT<u>C</u>A versus RT<u>C</u>A enrichments per sample within non-ecDNA *kataegis* (*top*) and non-SV associated *kataegis* (*bottom*), where YT<u>C</u>A and RT<u>C</u>A enrichment is suggestive of APOBEC3A or APOBEC3B activity, respectively. Genic mutations were divided into transcribed (template strand) and coding mutations. The RT<u>C</u>A/YT<u>C</u>A fold enrichments were compared to the fold enrichments of non-clustered mutations (p-values calculated using a Mann-Whitney *U*-tests and corrected for multiple hypothesis testing using the Benjamini-Hochberg false discovery rate procedure).

**Extended Data Figure 5: Recurrent mutagenesis and functional effects of ecDNA *kataegis***.

***a*)** A single head squamous cell carcinoma genome depicting the overlap of *kataegis* with ecDNA regions displayed as a rainfall (*top left)* with a single zoomed ecDNA represented using a circos plot (*top right*). The outer track of the circos plot represents the reference genome of the ecDNA with proximal known cancer driver genes. The middle track reflects a circular rainfall plot where each dot represents the intra-mutational distance (IMD) around a single mutation colored based on the substitution change. The innermost track shows the average variant allele frequency (VAF) for each *kataegic* event. *Bottom*: Two smaller regions of the selected ecDNA including a single *kataegic* event within *EGFR* resulting in a missense mutation and a single *kataegic* event within an intergenic region with the average VAF per event (orange). ***b*)** *Kataegic* substitutions found on ecDNA and resulting in recurrent coding mutations within known cancer genes.

## SUPPLEMENTARY INFORMATION

**Supplementary Note 1. Additional analysis of clustered mutagenesis**. The note contains information about differentiating between *omikli* and *kataegic* events as well as information about novel clustered mutational signatures.

**Supplementary Tables 1-6. Activities of mutational signatures causing clustered somatic mutations**. Each table provides the numbers of clustered mutations contributed by each signature in each sample. Table 1 contains the activities of doublet-base substitution signatures. Table 2 the activities of signatures generating multi-base substitutions. Table 3 the activities of signatures generating *omikli* events. Table 4 the activities of signatures generating *kataegic* events. Table 5 the activities of signatures generating clustered indel events. Table 6 the activities of signatures generating other clustered substitutions.

**Supplementary Tables 7. Coding *kataegic* mutations found within extrachromosomal circular DNA (ecDNA)**. Table provides the set of coding mutations as annotated by ENSEMBL’s Variant Effect Predictor tool by reporting only the most severe consequence^51^.

