## Extended Data Figures 1-6 for "Comprehensive analysis of clustered mutations in cancer reveals recurrent APOBEC3 mutagenesis of ecDNA"

**Extended Data Figure 1: Identification and classification of clustered events.**

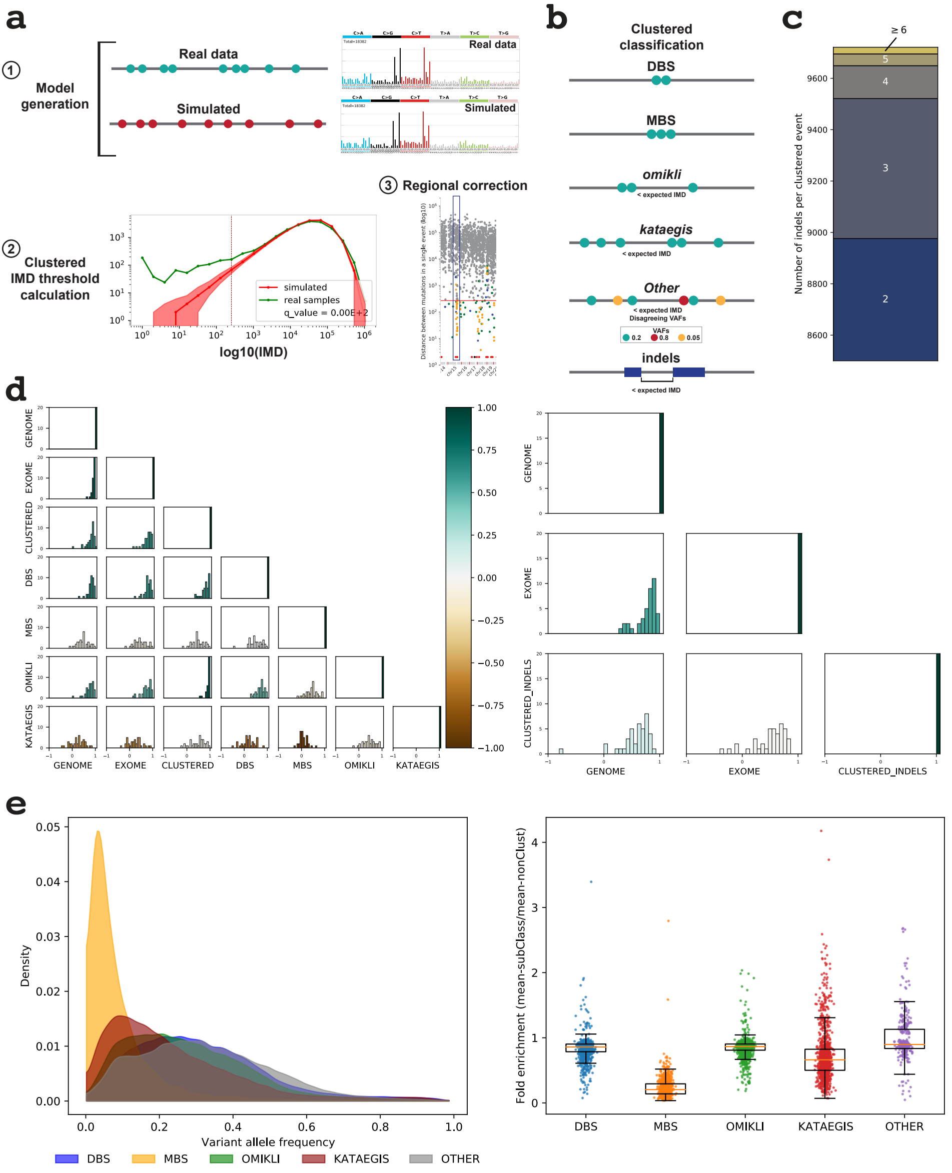

### Extended Data Figure 2: Experimental validation and epidemiological associations of clustered mutational processes

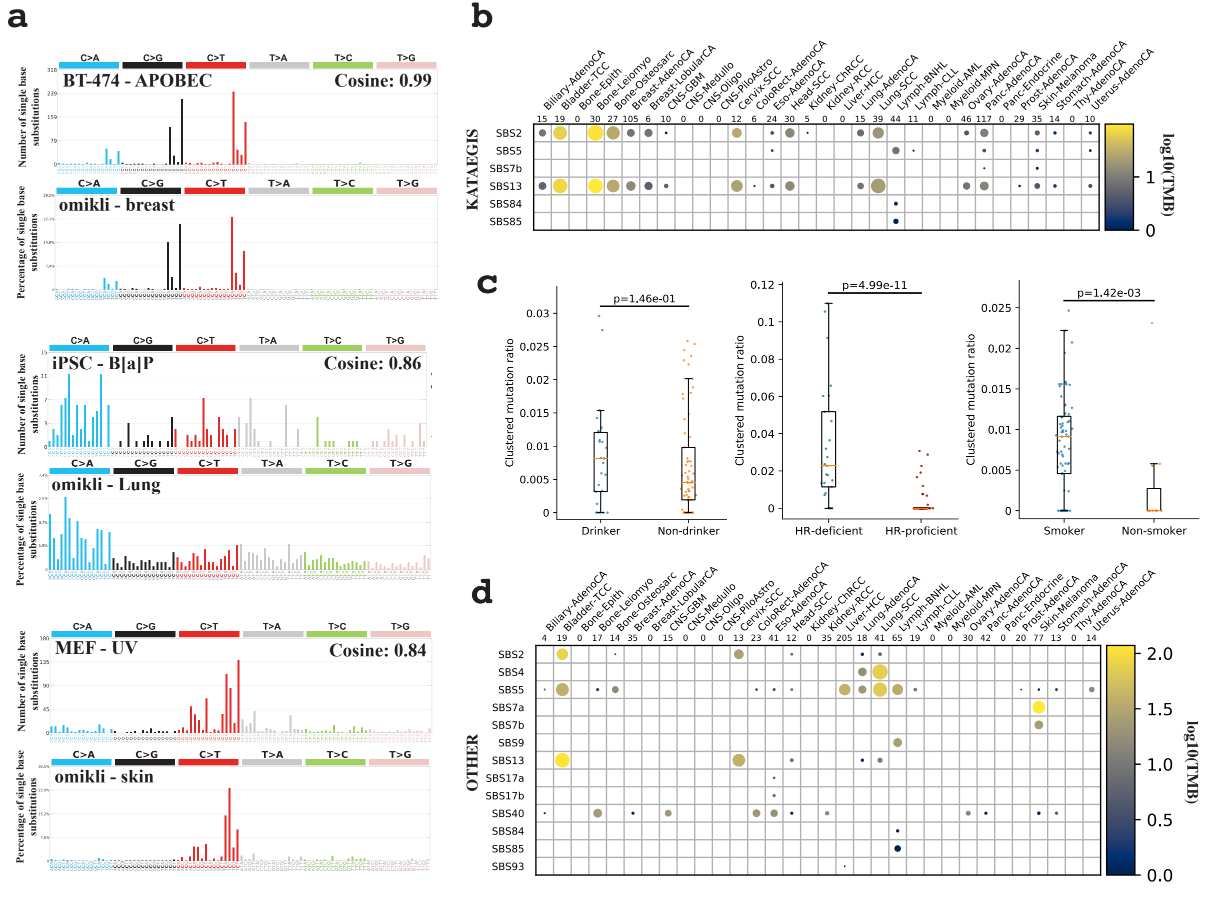

### Extended Data Figure 3: Mutational processes of clustered driver events

**a**

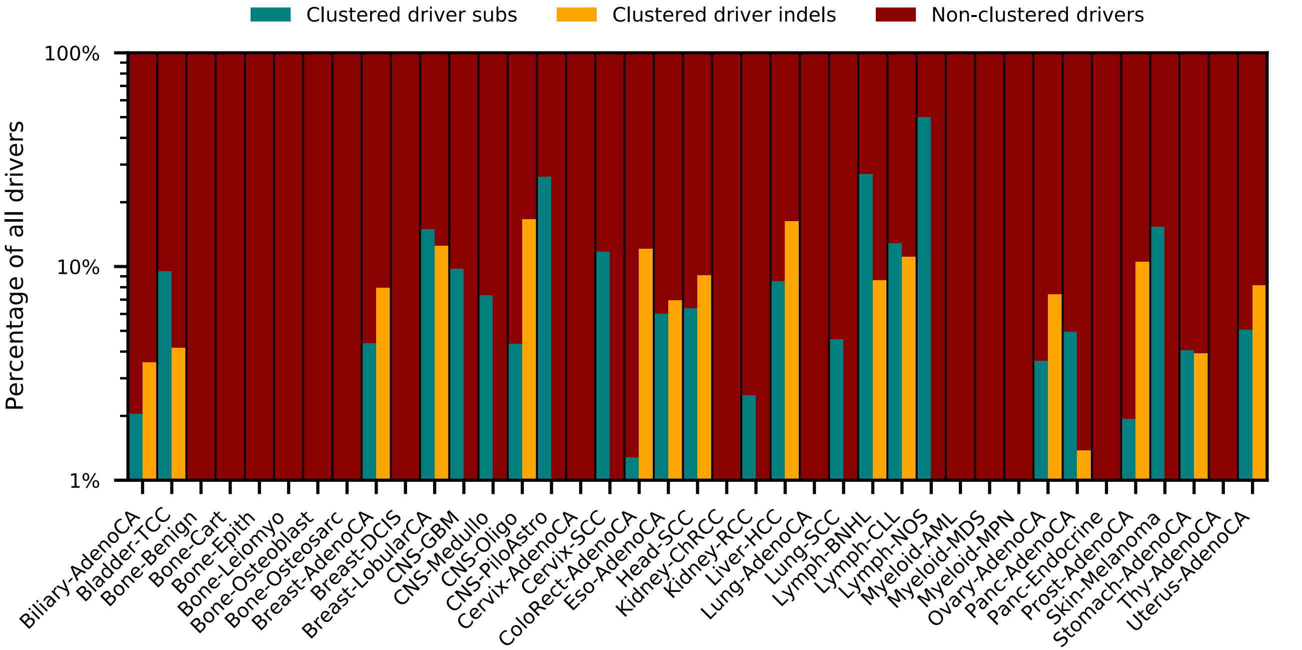


**b**

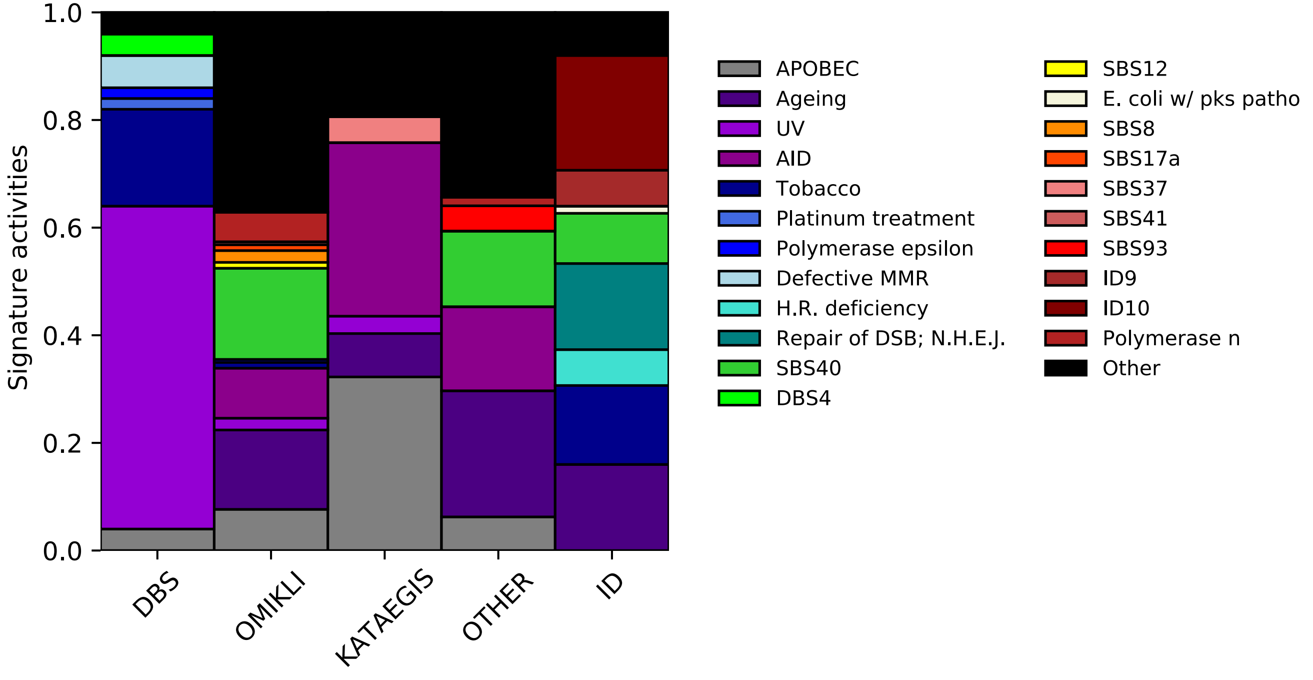

### Extended Data Figure 4: Clustered events and structural variations

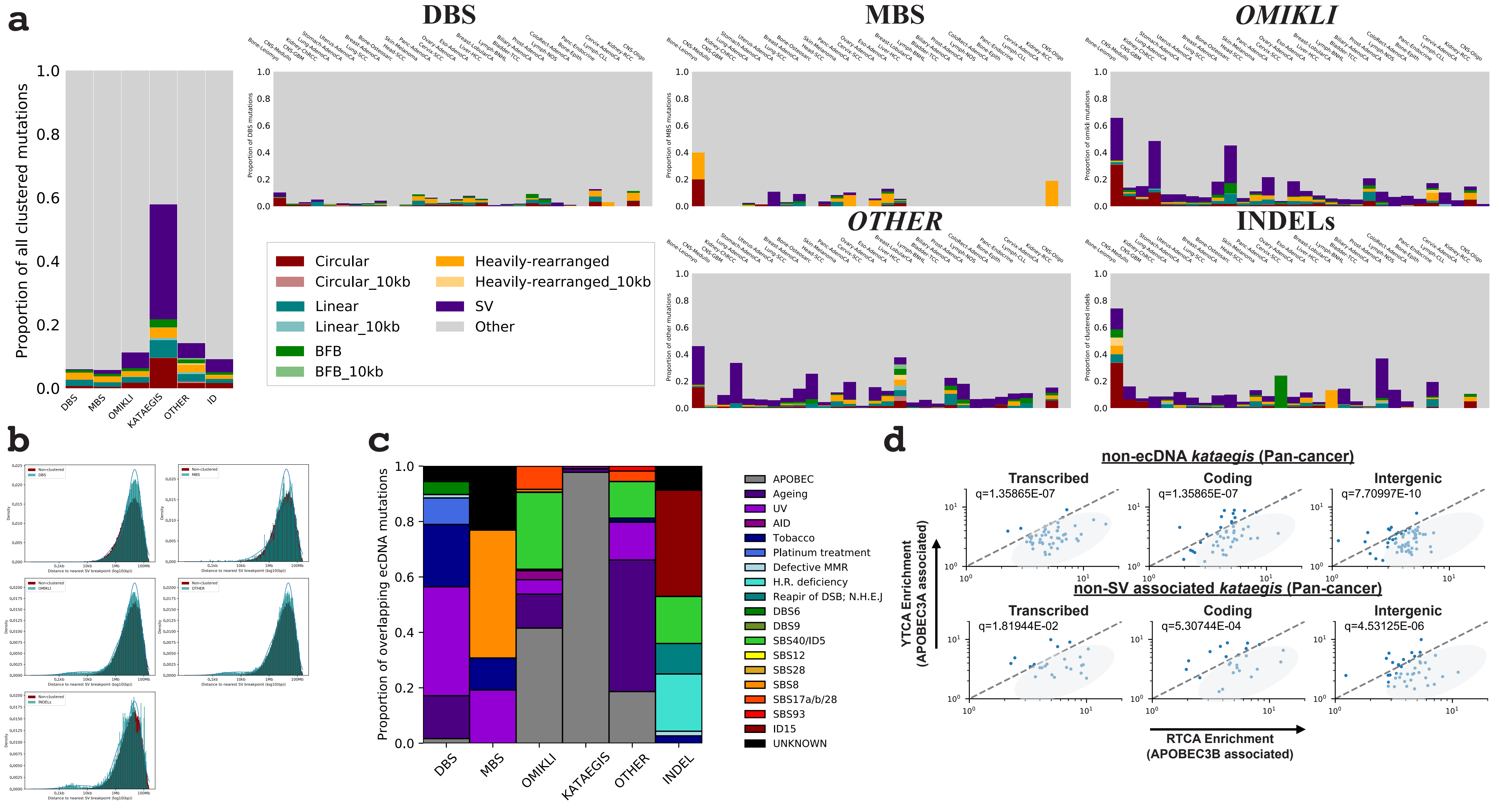

#### Extended Data Figure 5: Recurrent mutagenesis and functional effects of ecDNA *kataegis*

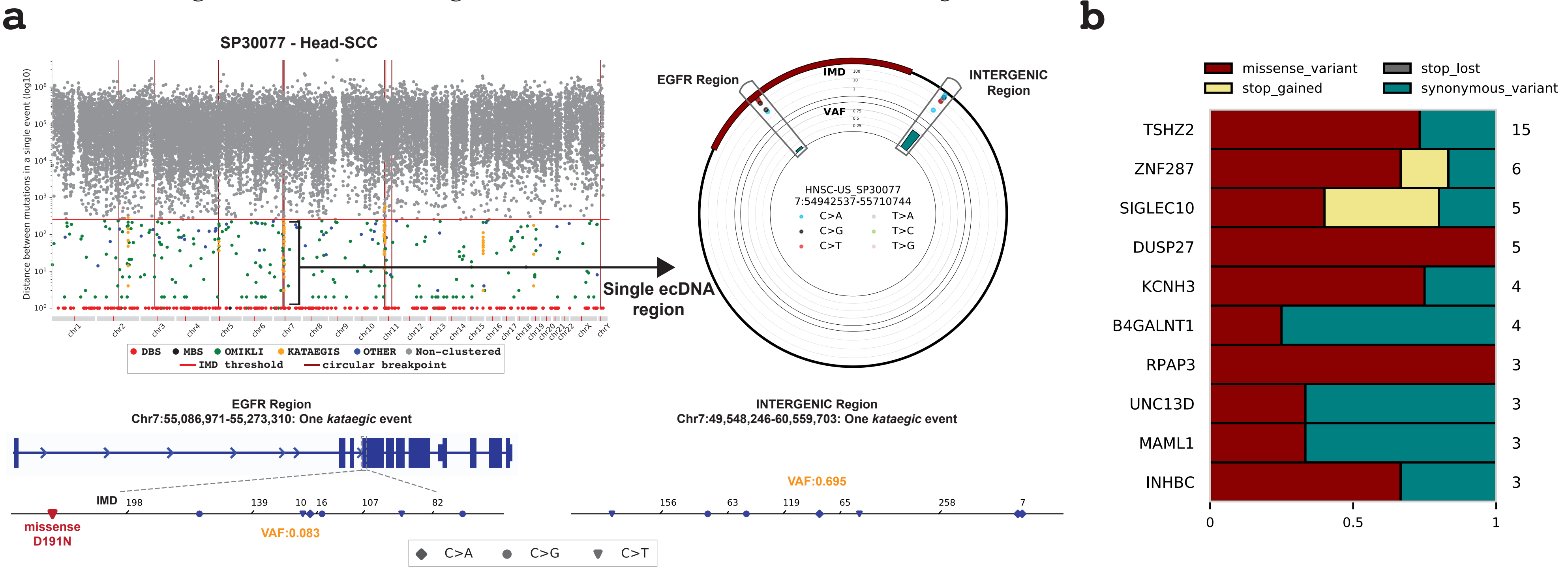
