## Supplementary Note 1 for "Comprehensive analysis of clustered mutations in cancer reveals recurrent APOBEC3 mutagenesis of ecDNA"

**Supplementary Note 1. Additional analysis of clustered mutagenesis.** The note contains information about differentiating between *omikli* and *kataegic* events as well as information about novel clustered mutational signatures.

### 1. Determining the number of clustered mutations in *omikli* and *kataegic* events.

To determine the cutoff of the number of mutations in an *omikli* versus a *kataegic* event, we modelled the distribution of clustered event sizes (excluding DBSs, MBSs, and *other* clustered events with disagreeable variant allele frequencies) using a mixture of two Poisson distributions (**Supplementary Figure 1*a***). The modelling also excluded clustered mutations from skin melanomas, that contribute a disproportionate number of DBS events, and clustered mutations from lymphomas, that contribute a large proportion of canonical and non-canonical AID *kataegis*. The first component, corresponding with *omikli* events (gold), had an average of 2.1 mutations per event, while the second component, corresponding to larger *kataegic* events (teal), had an average of 4.4 mutations per events. Using the posterior probabilities of each distribution, we calculated the likelihood of a given clustered event belonging to a specific component. Events comprised of four or more mutations were attributed to the *kataegic* component with >95% probability. Further, we assessed the IMD distributions of different sized events revealing approximately a 2-fold increase in average IMD between events possessing 3 and 4 mutations supporting the activity of two separate mutational processes (**Supplementary Figure 1*b***).

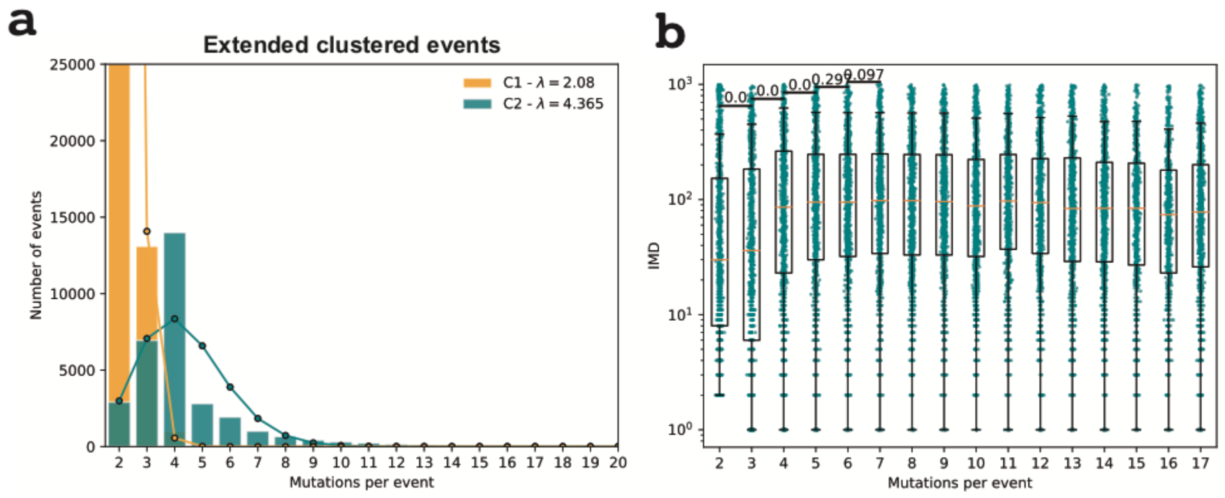


**Supplementary Figure 1: Determining the number of mutations differentiating between *omikli* and *kataegis*. *a)*** Modeling the number of mutations per event using a mixture of two Poisson distributions. The first component, representative of *omikli*, has an average IMD of 2.1, while the second component, representative of *kataegis*, has an average IMD of 4.4. The estimated contribution of mutations of each component are depicted as bars for each corresponding event size ***b)*** The distribution of IMDs per event across different sized events. The chosen cutoff between *omikli* and *kataegis* was four mutations.

2. **Identification of novel clustered mutational signatures.**

Our analysis of mutational signatures revealed 4 novel doublet-base substitution and 3 novel multi-base substitution signatures (**Supplementary Figure 2**). All of these were confined to individual cancer types and, in most cases, contributed mutations to a small number of samples.

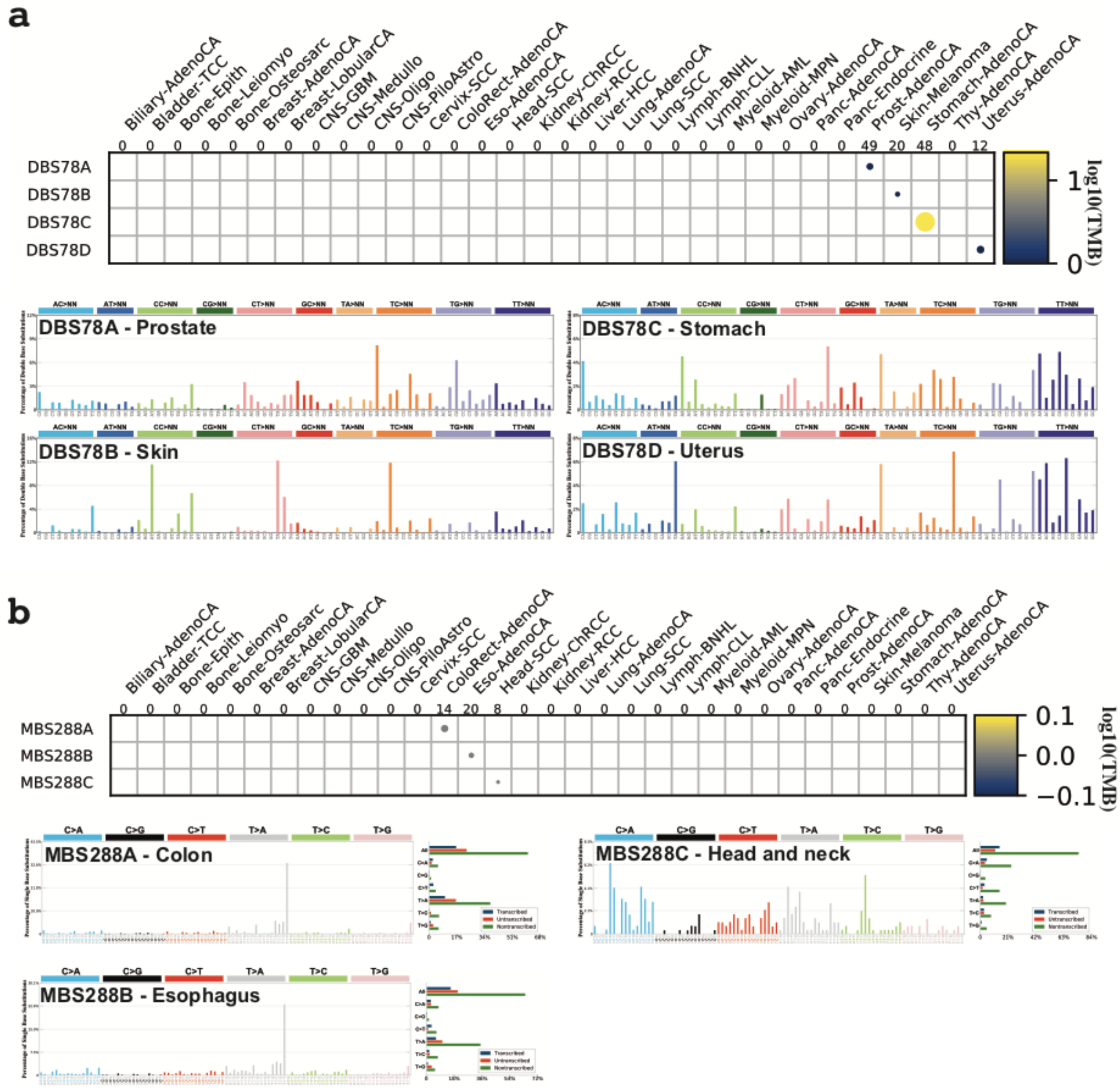


**Supplementary Figure 2:** ***De novo*** **signatures of doublet-base (DBS) and multi-base (MBS) signatures.** ***a)*** The activity of DBS *de novo* signatures (*top*) and the corresponding signatures extracted from prostate, skin, stomach, and uterine cancers that could not be accurately reconstructed using known COSMIC mutational signatures (*bottom*). ***b)*** The activity of MBS *de novo* signatures (*top*) and the corresponding signatures extracted from colon, esophagus, and head and neck cancers that could not be accurately reconstructed using known COSMIC mutational signatures (*bottom*).
